# Cells with Stochastically Increased Methyltransferase to Restriction Endonuclease Ratio Provide an Entry for Bacteriophage into Protected Cell Population

**DOI:** 10.1101/2022.03.28.486079

**Authors:** Alexander Kirillov, Natalia Morozova, Vasilisa Polinovaskaya, Sergey Smirnov, Mikhail Khodorkovskii, Lanying Zeng, Yaroslav Ispolatov, Konstantin Severinov

**Affiliations:** Skolkovo Institute of Science and Technology, Center for Life Sciences, Skolkovo, 121205, Russia; Peter the Great St Petersburg State Polytechnic University, St Petersburg 195251, Russia; Texas A&M University, Department of Biochemistry and Biophysics, Center for Phage Technology, College Station, TX 77843, USA; University of Santiago of Chile (USACH), Physics Department, Av. Víctor Jara 3493, Santiago, Chile; Waksman Institute for Microbiology and Department of Molecular Biology and Biochemistry, Rutgers, State University of New Jersey, Piscataway, NJ 08854, USA

**Keywords:** restriction-modification system, bacteriophage infection, plasmid copy number

## Abstract

The action of type II restriction-modification (RM) systems depends on restriction endonuclease (REase), which cleaves foreign DNA at specific sites, and methyltransferase (MTase), which protects host genome from restriction by methylating the same sites. We show that protection from phage infection increases as the copy number of plasmids carrying the Esp1396I RM system is increased. However, since increased plasmid copy number leads to both increased absolute intracellular REase and MTase levels and decreased MTase to REase ratio, it is impossible to determine which factor determines resistance/susceptibility to infection. By controlled expression of Esp1396I MTase or REase genes in cells carrying the Esp1396I system, we show that a shift in the MTase to REase ratio caused by overproduction of MTase or REase leads, respectively, to decreased or increased protection from infection. Consistently, due to stochastic variation of MTase and REase amount in individual cells, bacterial cells that are productively infected by bacteriophage have significantly higher MTase to REase ratios than cells that ward off the infection. Our results suggest that cells with transiently increased MTase to REase ratio at the time of infection serve as entry points for unmodified phage DNA into protected bacterial populations.

## INTRODUCTION

In nature, bacteria are constantly under attack from viruses (bacteriophages) (Hatfull and Hendrix, 2011). Multiple mechanisms of bacterial defense against viral infection have evolved. One common resistance mechanism is provided by type II restriction-modification (RM) systems (Pingoud et al., 2014, Makarova et al., 2013, Vasu and Nagaraja, 2013). The defensive action of these systems is due to activity of two enzymes, a restriction endonuclease (REase) and a cognate methyltransferase (MTase). Both enzymes recognize identical, usually 4-6 bp, sites in the DNA. An REase cleaves (restricts) DNA at or close to unmodified sites, while the cognate MTase methylates (modifies) specific bases in the recognized sites. Because most recognition sites are palindromic, MTases introduce two methyl groups at both strands of the recognized sequence, generating first hemimethylated and then fully methylated sites, neither of which are recognized by cognate REases and are thus not subject to restriction (Pingoud et al., 2005, Roberts et al., 2003, Tock and Dryden, 2005). The RM systems clearly impact on phages, which respond to this pressure in multiple ways including avoidance of RM recognition sites in their genomes, encoding special anti-restriction proteins, heavily modifying bases of their DNA, or substituting thymine residues for uracils (Rusinov et al., 2018, Spoerel et al., 1979, Serfiotis-Mitsa et al., 2010, Hutinet et al., 2019, Samson et al., 2013, Stern and Sorek, 2011).

While the presence of an RM system in a bacterial host provides strong protection from a viral attack (decreasing the efficiency of infection in common viral plaque-forming assays by as much as million-fold compared to unprotected hosts), this protection is short-lived (Luria and Human, 1952). Progeny phages with modified DNA invariably arise at least at conditions of laboratory experiments. Once such epigenetically modified phages appear, they productively infect cells with a cognate RM system, rendering the defense useless. On the other hand, when the modified phage infects a cell without an RM system, unmodified phage progeny is released and the initial level of protection is restored.

The intracellular amounts/activity levels of the MTase and REase enzymes shall be tightly controlled because a disbalance can either lead to the death of uninfected (excess REase activity) or infected (excess MTase activity) cells (Ichige and Kobayashi, 2005). Special mechanisms have evolved to allow highly controlled expression of genes coding for RM enzymes (Mruk and Kobayashi, 2014). These mechanisms act through multiple positive and negative feedback loops and ensure homeostasis of RM gene expression. Yet, the fact that RM protection can be overcome with appreciable (∼10^−3^) probability suggests that in some infected cells at least hemimethylation of each of the recognition sites in the phage genome must occur before even a single restriction event takes place. In principle, this can happen when activities of the MTase and REase enzymes in an infected cell are altered such that excessive MTase activity allows preemptive modification of phage DNA. In contrast, excessive REase activity should allow better protection from the phage but may lead to cleavage of bacterial DNA in the absence of infection.

Earlier, we used mathematical modeling to show how the ability of a phage to overcome protection afforded by an RM system can be determined by the relative ratio of MTase and REase activities in the cell (Enikeeva et al., 2010). To verify the model predictions we here experimentally perturbed the intracellular amounts of MTase and REase of the Esp1396I RM system (Cesnaviciene et al., 2003, Bogdanova et al., 2009). Similarly to many other type II RM systems, expression of Esp1396I MTase and REase genes is tightly regulated by a transcription factor (C-protein) that is part of the Esp1396I gene cluster. Through multiple interactions with operator sites located in front of the Esp1396I MTase and REase genes as well as its own gene, the Esp1396I C-protein finely controls the steady-state amounts of Esp1396I MTase and REase present inside the cell (Bogdanova et al., 2009) and allows coordinated synthesis of both enzymes when a plasmid containing the Esp1396I RM system enters a naive cell (Morozova et al., 2016). In this work we determined the effects of increasing the amounts of Esp1396I enzymes and of their ratios on efficiency of phage infection and the half-life of unmodified phage DNA inside the cells. In agreement with earlier theoretical analysis, we show that changing the ratio of MTase and REase enzymes while keeping the intracellular concentration of one of the enzymes constant can have a drastic effect on the level of protection from phage infection. By examining the levels of stochastic variation of MTase and REase enzymes’ amounts in individual cells before and after phage infection, we show that cells with increased MTase to REase ratio at the time of infection serve as entry points for unmodified phage into protected bacterial populations.

## MATERIALS AND METHODS

### Bacterial strains and plasmids

*E. coli* DH5*α* strain was used for cloning, flow cytometry and most microscopy experiments. *E. coli* K-12 MG1655 cells (dam-, SeqA protein is fused with the mKO2 orange fluorescent protein) were used to observe and measure bacteriophage λ_vir_ DNA degradation rate. *E. coli* Rosetta strain was used for fluorescent proteins (Venus and mCherry) expression from pET-based plasmids. Cells were grown in standard LB medium (Amresco) with antibiotics and inducers (if necessary). Antibiotics were added at concentrations of 100 *μ*g/ml for ampicillin and 25 *μ*g/ml for chloramphenicol. Plasmid transformation was performed using appropriate ultra-competent cells (Inoue et al., 1990). Expression of Venus and mCherry proteins from pET-based plasmids was performed by the addition of 1 mM IPTG to cell cultures. Expression of MTase or REase from *araBAD* promoter was performed by the addition of different concentrations (from 1 *μ*M to 12 mM) of L-arabinose (Ara) to cell cultures. To prevent leaky expression from the *araBAD* promoter (if needed), glucose at concentrations of 0.2 % was added.

pET21a-based expression plasmids with Venus or mCherry genes and the fluorescently-labeled Esp1396I*_*fluo RM system were described previously (Morozova et al., 2016). Additional Esp1396I_fluo plasmids (pRM-Low, pRM-Mid and pRM-High) and plasmids expressing separate Esp1396I REase::mCherry gene or Esp1396I MTase::Venus gene fusions (pRM-Mid+R and pRM-Mid+M) were constructed using standard molecular cloning techniques. Details are available from the authors upon request.

### Phage protection plaque assay

Plaque assay was performed using DH5α strain carrying empty pLow, pMid, or pHigh plasmids (unprotected cells); protected cells carrying the pRM-Low, pRM-Mid, or pRM-High plasmids; and cells carrying the pRM-Mid and +R or +M plasmids with induction with Ara. The overnight cultures were grown in LB with the addition of ampicillin (pLow, pMid, pHigh, pRM-Low, pRM-Mid, or pRM-High) or ampicillin and chloramphenicol (pRM-Mid+R or +M) at 37°C. Cells were diluted 1:100 in fresh LB medium with the addition of 10 mM MgSO_4_, 0.2% maltose and antibiotics and grown at 37°C to OD_600_ ≈ 0.4. Petri dishes were precast with 10 ml of 1.5% LB agar (bottom layer) and then overlaid with 4 ml of 0.4% LB agar (top layer) mixed with the 100 μl of cell cultures, 100 μl of serial dilutions of bacteriophage lysate, 10 mM MgSO_4_, 0.2% maltose, and appropriate antibiotics. The plates were incubated for 10 hours at 37°C and phage titers were determined as described in (Mazzocco et al., 2009).

### Protein purification

Purified Venus and mCherry fluorescent proteins were used as standards for microscopy calibration (Morozova et al., 2016). The Rosetta strain was transformed with the pET21a-based Venus or mCherry expression plasmids. Overnight cultures from the single colonies were grown in LB medium with the addition of ampicillin and chloramphenicol at 37°C. Cells were diluted 1:100 in 50 ml of fresh LB medium with the addition of ampicillin and chloramphenicol and were grown at 37°C till OD_600_ = 0.6. Protein expression was induced by the addition of 1 mM IPTG. After 4 hours of growth, cells were centrifuged at 4000 g, and the pellet was resuspended in 50 mM Tris-HCl, 500 mM NaCl, 10 mM imidazole. Cells were lysed by sonication in the presence of lysozyme (1 mg/ml) and PMSF (1 mM). The Venus and mCherry proteins C-terminally tagged with a 6-His-tag were affinity-purified from the cleared cell lysate on a 1-ml Ni-NTA agarose column (Crowe et al., 1994). After the loading of cell lysate onto the column, it was washed with three volumes of 50 mM Tris-HCl, 500 mM NaCl, 10 mM imidazole buffer and protein was eluted using 50 mM Tris-HCl, 500 mM NaCl, 400 mM imidazole buffer. Fractions containing purified target proteins (∼95% pure as judged by visual inspection of SDS gels) were diluted with glycerol (50% final concentration), frozen in liquid nitrogen, and stored at -80°C prior to use. The concentration of proteins was measured using the Bradford assay (Bradford, 1976).

### Fluorescence intensity calibration

Purified Venus and mCherry proteins with known concentrations were used as a standard for spectrofluorimetric (Varian Cary Eclipse) measurements from which the fluorescence of a single fluorescent protein molecule was calculated. Fresh cultures of *E. coli* Rosetta cells with pET-based Venus or mCherry expression plasmids were grown in LB medium with the addition of ampicillin and chloramphenicol and 1 mM IPTG. The concentration of cells was measured using live microscopy, and culture fluorescence was measured using Varian Cary Eclipse fluorescence spectrophotometer. This data allowed calculating the average fluorescence of a single cell and, consequently, the average number of fluorescent proteins per cell. Simultaneously, average fluorescence of a single cell observed by live microscopy was measured using ImageJ (Fiji) tools (Schindelin et al., 2012). The autofluorescence signal was calculated from data obtained with microscopy of cells harboring empty plasmid vectors. After subtraction of mean autofluorescence signal, the fluorescence of a single molecule fluorescent protein molecule under the microscope was calculated. This allowed us to determine the value of a coefficient for recalculation of fluorescence intensity of a cell to the number of fluorescent protein molecules present in it.

### Microscopy tools

Fluorescence microscopy was performed using a Nikon Eclipse Ti-E inverted microscope equipped with a custom incubation system. Fluorescence signals in Venus and mCherry fluorescence channels were detected using Semrock filter sets YFP-2427B and TxRed-4040C, respectively. The long-time microscope filming was performed using Micro-Manager 2.0 tools (Edelstein et al., 2014). Time-lapse filming was performed using sCMOS ZYLA 4.2MP Plus Andor camera. Image analysis was performed using ImageJ (Fiji) (Schindelin et al., 2012) with the Circle4px plugin used for measurements of fluorescence in individual cells.

### Microscopy Slide preparation

The double-sided tape (Scotch) was used for the frame formation on one microscopic Slide. The 1.5% agarose (Helicon) diluted in 0.25% LB (Amresco) was used to make a smooth surface inside the frame. The agarose pad was pressed by the second similar Slide. After agarose solidification, the upper Slide was removed. Thin agarose strips were cut to provide oxygen to the sample in the chamber. The sample mixture was put on the thin agarose pads. When the mixture was fully absorbed, the chamber was sealed with the coverslip (24 × 40 mm, Menzel-Gläser).

### Microscopy of phage-infected cells

The fresh culture of *E. coli* DH5*α* cells carrying the pRM-Low plasmid was grown to OD_600_ = 0.4 in LB supplemented with 10 mM MgSO_4_, 0.2% maltose and ampicillin. Cell culture was incubated at 37°C for 5 minutes with the phage (MOI=1) to allow adsorption. Microscope Slides were prepared as described above and 1 μl of cell culture mixed with phage was added on the agarose pad. The filming was carried out in three different channels: transmitted light (TL), Venus fluorescence channel, and mCherry fluorescence channel. The level of autofluorescence was measured using *E. coli* DH5*α* cells carrying the pLow plasmid. Each field of view was imaged every 10 minutes for 300 minutes.

### Observation of bacteriophage λ _vir_ DNA inside bacterial cells

The method is described in detail elsewhere (Shao et al., 2015). The culture of cells with Dam MTase gene deletion (*E. coli* K-12 MG1655) was grown at M9 to OD_600_ = 0.4 with the 10 mM MgSO_4_ and 0.2% maltose. The cells were mixed with bacteriophage λ_vir_ whose DNA is fully-methylated by Dam MTase and incubated for 5 minutes. A 1 μl of the mixture was placed on an agarose pad containing M9 medium. The time-lapse filming was performed using a Nikon Eclipse Ti-E inverted microscope. The filming was carried out in transmitted light (TL) and mKO2 fluorescence channels for the bacteriophage DNA visualization. Visualization was performed due to the detection of SeqA::mKO2 protein which specifically binds methylated or hemimethylated by Dam MTase DNA. The distinct foci were visualized using the z-stacks. Each field of view was imaged every 15 minutes for more than 150 minutes.

### Sample preparation for flow cytometry analysis

Fresh culture of *E. coli* DH5*α* cells transformed with the pRM-Low plasmid was grown to OD_600_ = 0.4 in 10 ml of LB with ampicillin. 5 ml of cell culture was mixed with λ_vir_ phage lysate at the MOI=5. The remaining uninfected cells were used as control. Using microscopy, we determined that the end of the first infection cycle in our conditions occurred ∼1 hour after the addition of the phage (data not shown). Thus, cells were precipitated by centrifugation 55 minutes after the infection, when productively infected cells were already lysed, while the secondary infection have either not yet started or were at the very beginning. The supernatant was discarded, and the cell pellet was resuspended in phosphate-buffered saline (PBS) to reach a ∼10^6^ bacteria/ml concentration.

### Flow cytometry data analysis

FACS Aria III (BD) was used for fluorescence measurements. The area of bacterial cells was gated by comparing the forward and side scatter plots of PBS and plasmid-free *E. coli* DH5α cells resuspended in PBS. The 488-nm and 561-nm lasers were used for fluorescence excitation in green (Venus) and red (mCherry) channels, respectively. The Blue 530/30-A and Yel-Green 610/20-A filters were used for fluorescence detection. The selection of the region corresponding to bacterial cells was chosen by comparing the control and cell-containing samples on the side-forward scatter (SS-FS) plots. 20,000 events (fluorescence intensities of individual cells) were recorded for each sample of cells carrying the pRM-Low plasmid without and after a round of λ_vir_ infection. The obtained data were processed using Weasel and CytExpert software and the «R» programming language (R Core Team, 2013). Samples with and without viral infection were analyzed using the Kolmogorov-Smirnov test (Young, 1977). The null hypothesis was the assumption that the distributions with and without infection do not differ.

## RESULTS

### The copy number of plasmids carrying an RM system affects the level of protection from phage infection

Three plasmids carrying the Esp1396I restriction-modification system were created: pSC10l_Esp1396I_fluo, pBR322_Esp1396I_fluo, and pUC19_Esp1396I_fluo. The copy numbers (PCN) for the corresponding plasmid vectors were reported to be about 5, about 20, and more than 100 copies per cell, respectively (Jahn et al., 2016), and are consistent with estimates obtained using real-time PCR of total DNA samples from cells carrying Esp1396I plasmids (Suppl. Figure S1). Heretofore, we will refer to pSC101_Esp1396I_fluo, pBR322_Esp1396I_fluo, and pUC19_Esp1396I_fluo as pRM-Low, pRM-Mid, and pRM-High plasmids, correspondingly. The Esp1396I system cloned on each plasmid is modified by fusing the MTase and REase genes to Venus and mCherry fluorescent proteins genes, respectively. Elsewhere we show that such a modified system is functional and that both fusion proteins remain intact within the cell (Morozova et al., 2016).

Figure 1(a) shows microscopy images of live cells carrying fluorescently-labeled Esp1396I RM system on different plasmids in Venus and mCherry fluorescence channels. As a control, cells with an empty vector pBR322 (pMid) were used. Overall, higher fluorescence (and, therefore, higher levels of fusion proteins) were observed in cells with higher-copy number plasmids. Compared to cells carrying the pRM-High plasmid, the fluorescence intensity of cells carrying the pRM-Mid plasmid was decreased 2.6-fold in the mCherry channel and 1.4-fold in the Venus channel. In cells carrying the pRM-Low plasmid the decrease was more pronounced (9-fold and 2.1-fold respectively) (Figure 1(a), (b)). Since one molecule of MTase or REase corresponds to one molecule of Venus or mCherry proteins, respectively, the number of Esp1396I enzymes’ molecules and their ratios in individual cells could be estimated from fluorescence intensity in individual cells (Figure 1(d), (e), (f)). As can be seen, while the absolute number of RM enzymes was proportional to PCN, as expected, the dependence of MTase and REase amounts on PCN was not the same. The correlation coefficients between REase::mCherry and MTase::Venus concentrations in individual cells were 0.48 (n = 666) in cells with pRM-High, 0.45 (n = 690) in cells with pRM-Mid, and 0.37 (n = 721) in cells with the pRM-Low plasmid. The differences in correlation coefficients can be explained by the different widths of MTase to REase ratio distributions in individual cells, which were wider in cells with lower copy number plasmid (Figure 1(f)). The mean MTase to REase ratio values increased with decreasing PCN. While the concentration of the Esp1396I C-protein that regulates transcription of the MTase to REase genes was not measured, it is possible that the observed effects are caused by lower C-protein concentrations in cells carrying lower copy number plasmids, which should lead to higher relative MTase expression levels and larger stochastic effects (Bogdanova et al., 2009).

**Figure 1.**
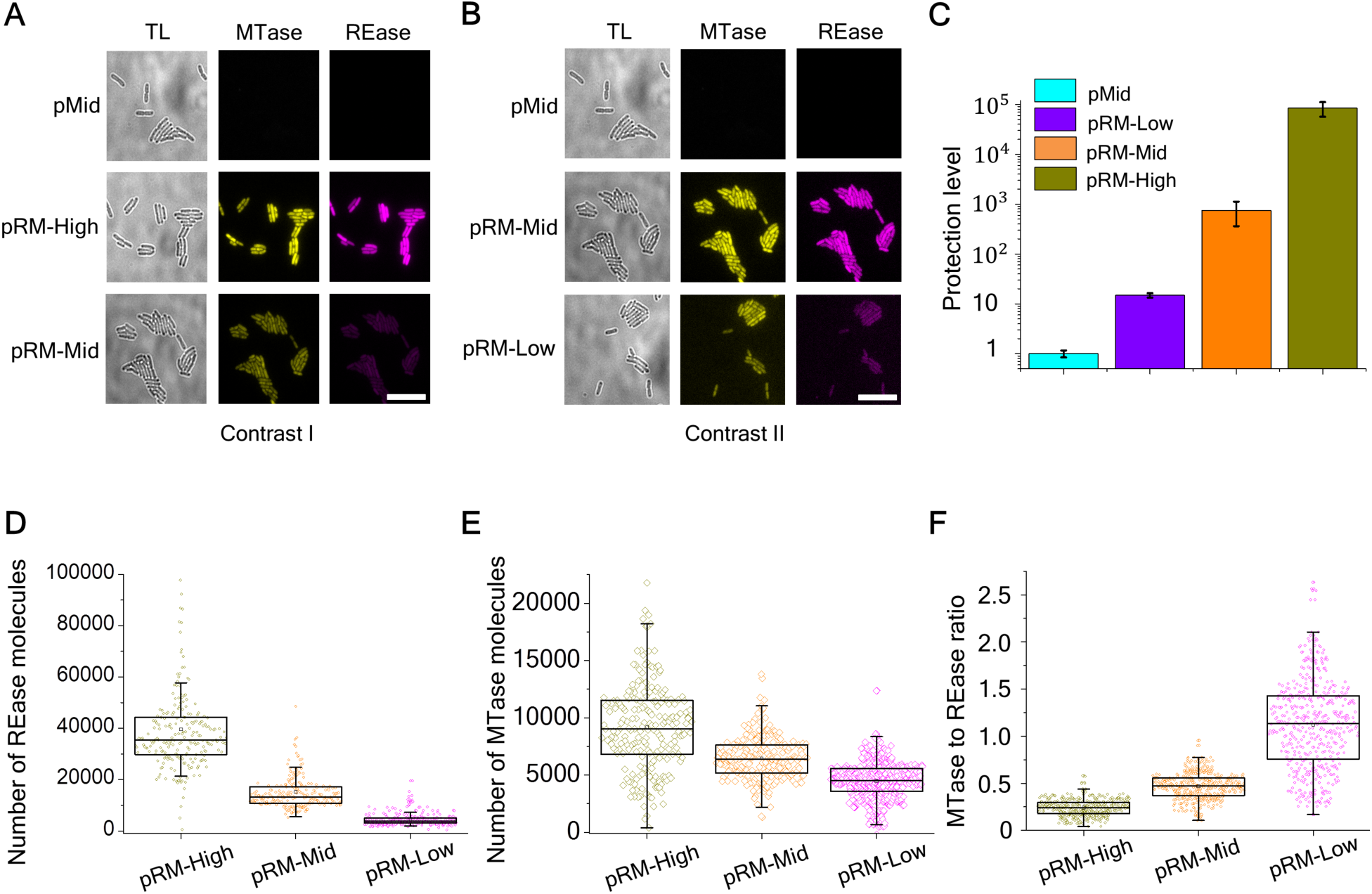
Effects of RM plasmid copy number on intracellular concentrations of Esp1396I enzymes and levels of protection from phage infection. (A and B) Images of cells with an empty pBR322 plasmid vector (“pMid”), and cells transformed with indicated Esp1396I_fluo plasmids are shown in transmitted light (TL), Venus (yellow), and mCherry (magenta) channels. Scale bar is 20 μm. The contrasts in (A) and (B) are different to reveal the differences in fluorescence signals. (C) Protection levels of cells carrying indicated RM plasmids or the pMid vector from phage λ_vir_ infection are shown. Protection levels are calculated by dividing the titer of phage lysate on the lawn of cells carrying the pMid vector by a titer determined on lawns of cells carrying an indicated plasmid. Bars represent mean protection levels obtained from three independent experiments. Error bars show standard errors of the mean. (D)-(F) The distributions of REase (D) and MTase (E) molecule numbers and of MTase to REase ratios (F) in individual cells carrying the Esp1396I_fluo system on indicated plasmids.

Protection levels of cells carrying different RM plasmids or empty vector controls were tested by applying drops of serial dilutions of phage λ_vir_ lysate on the surfaces of lawns formed by different cells. The level of protection was calculated as a ratio of phage titers determined on lawns of cells with RM plasmids to the titer observed on control cell lawns. As expected, no difference in resistance was observed between control cells carrying different empty vectors (Suppl. Figure S2). As can be seen (Figure 1(c)), bacteria harboring the pRM-Low plasmid were least protected (∼10-fold decrease in phage titer), while bacteria carrying the pRM-High plasmid were most protected (∼10^5^-fold phage titer decrease). Cells carrying the pRM-Mid plasmid had an intermediate (∼10^2^-10^3^-fold) protection level. We conclude that there is a positive correlation between the copy number of an RM system-encoding plasmid and the protection level from phage infection. Despite low protection level of cultures carrying the pRM-Low plasmid, the progeny phage formed upon a single-round of infection of these cells productively infected the highly protected pRM-High cultures and hence was modified (Suppl. Figure S3).

### The influence of additional REase or MTase enzymes on protection from bacteriophage infection

Our results show that the higher the copy number of a plasmid carrying the Esp1396I system, the higher is the average concentration of both RM enzymes in the cells and the level of protection against viral infection. However, when the copy number of a plasmid carrying the RM system is increased, the relative concentration of REase increases to a higher extent than that of the MTase, resulting in a decrease of the MTase to REase ratio. To find out how changes in the concentration of only one RM enzyme would affect protection against the virus, gene fusions encoding the REase::mCherry or the MTase::Venus proteins were placed downstream of the arabinose-inducible promoter on pACYC plasmids compatible with the pRM-Mid plasmid. We will refer to the resulting pACYC_R_fluo and pACYC_M_fluo plasmids as “+R” and “+M”, respectively. Bacteria co-transformed with pRM-Mid and either +R or +M plasmids were analyzed by fluorescence microscopy in the presence or in the absence of 10 mM arabinose inducer. As a control, cells without the RM system plasmid and cells that carried the pRM-Mid plasmid and the empty pACYC vector were used. As can be seen from Figure 2(a), induction of the MTase gene expression led to increased fluorescence in the Venus channel compared to control bacteria containing the pRM-Mid plasmid and the empty vector, as expected. Also, as expected, an increase in fluorescence in the mCherry channel was observed upon induction of additional REase synthesis.

**Figure 2.**
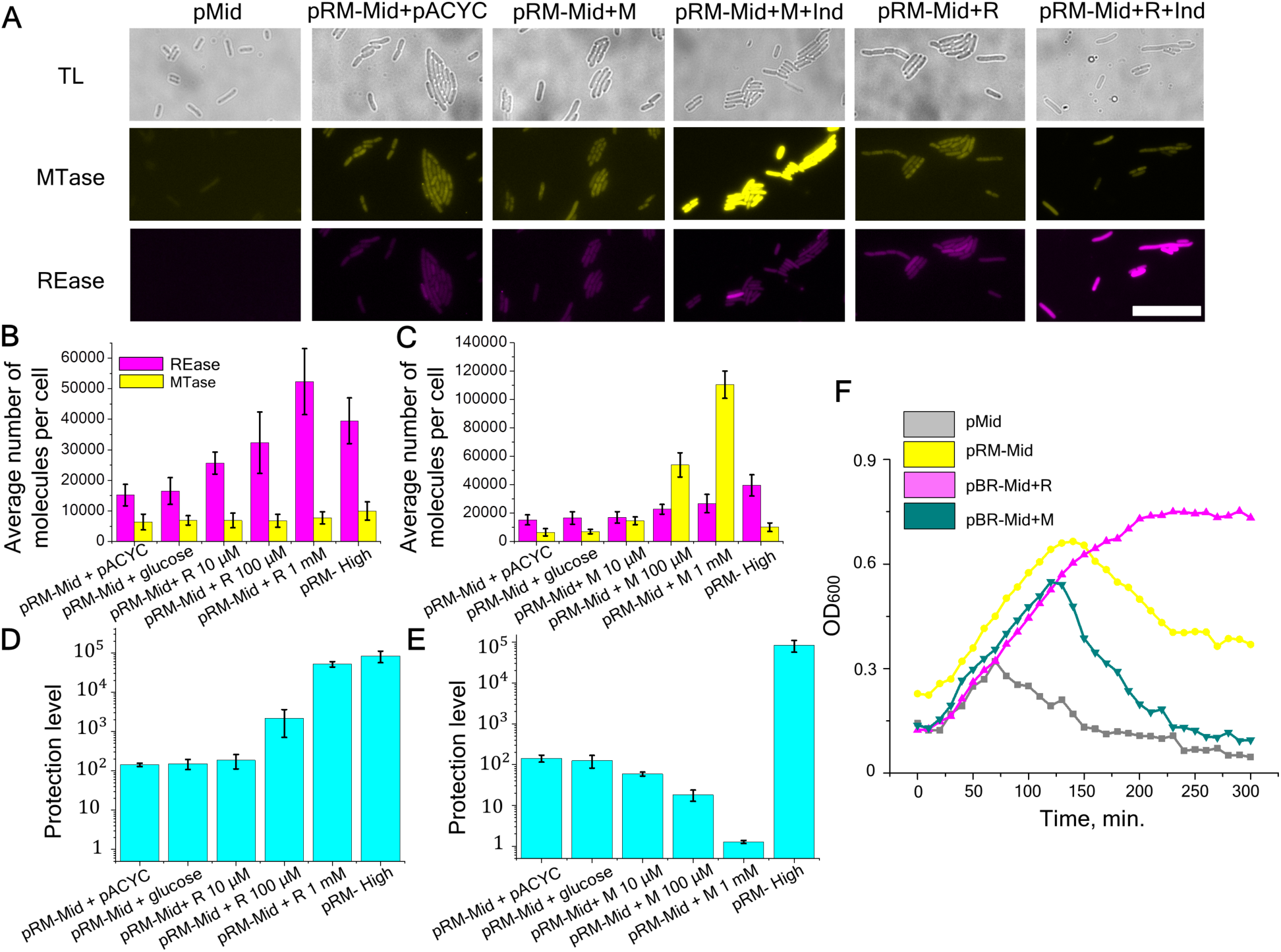
The influence of additionally induced Esp1396I enzymes on the level of protection. (A) Images of cells in transmitted light (gray) and in two fluorescence channels Venus (yellow) and mCherry (magenta): pMid -control cells without the RM system; pRM-Mid + pACYC - cells with pRM-Mid and pACYC plasmids; pRM-Mid+M - cells with pRM-Mid and +M plasmids with or without 10 mM arabinose induction; pRM-Mid+R - cells with pRM-Mid and +R plasmids with or without 10 mM arabinose induction). Scale bar is 20 μm. (B and C) Intensities of fluorescence in 300 individual cells with additional synthesis of REase (B) or MTase (C) induced by different concentrations of arabinose. Yellow and magenta bars represent the mean number of MTase and REase molecules per cell, respectively. Error bars represent standard error of the mean. (D and E) Protection from phage infection of cells with additionally expressed REase (D) or MTase (E). Protection levels were determined as described in Figure 1(c) legend. Error bars represent standard error of the mean. (F) Growth of cultures of cells without RM protection (pMid), cells carrying the pRM-Mid plasmid, and pRM-Mid carrying cells expressing additional MTase (pRM-Mid+M) or REase (pRM-Mid+R). Cells were grown in the presence of 1 mM arabinose. At zero time point, cultures were infected with λ_vir_ phage at a MOI of 0.2.

The increase in REase or MTase production in cells carrying additional expression plasmids as a function of arabinose inducer concentration was measured 2 hours post-induction using fluorescence microscopy. As controls, we used *i*) cells carrying empty pMid and pACYC vectors; *ii*) cells with the pRM-Mid plasmid and the pACYC vector, and *iii*) cells with the pRM-Mid plasmid and inducible REase or MTase expression plasmids grown in the presence of 0.2% glucose, a condition that should completely repress transcription from the arabinose promoter. Mean fluorescence values for 300 cells from each sample were determined and results are presented in Figure 2(b). For cells harboring the inducible REase fusion, induction with 0.01 and 0.1 mM arabinose led to, respectively, 1.5-and 1.7-fold increase of mean fluorescence intensity in the mCherry channel compared to cells with the pRM-Mid plasmid and the pACYC vector or cells grown in the presence of glucose. A 2.7-fold increase was detected in cells induced with 1 mM arabinose. Further increase of inducer concentration did not lead to additional increase in mean fluorescence intensity (data not shown). No increase in mean fluorescence in the Venus channel in samples with additionally induced REase was observed, indicating that levels of MTase remain constant. In cultures of cells with inducible MTase, mean fluorescence intensity in the Venus channel in the presence of 0.01, 0.1, and 1 mM of arabinose increased 1.8-, 8.7-, and 18-fold, respectively (Figure 2(c)). In the presence of 0.1, and 1 mM arabinose, a moderate (1.6-1.7 fold) increase in the level of REase was also detected. Overall, we conclude that the use of additional plasmids expressing REase or MTase allows us to alter the ratio of RM enzymes without significantly affecting the concentration of the counterpart protein expressed from the RM system-carrying plasmid. The concentration of REase in cells harboring the pRM-Mid plasmid and expressing additional REase in the presence of 0.1-1 mM arabinose approaches that in cells harboring the pRM-High plasmid (Figure 2(b)). The concentration of MTase in cells harboring the pRM-Mid plasmid and expressing additional MTase at 1 mM arabinose is several-fold higher than that in cells harboring the pRM-High plasmid (Figure 2(c)).

We next determined the level of protection for cells with different MTase to REase ratios. In samples with inducible REase, the presence of 0.01, 0.1, and 1 mM arabinose increased the level of protection by 0.6, 1, and 2 orders of magnitude, respectively, compared to that of cells carrying the pRM-Mid plasmid and the pACYC vector (Figure 2(d)). At 1 mM arabinose, the protection was equal to that observed in cells harboring the pRM-High plasmid. The increase of inducer concentration to 10 mM did not lead to additional protection (data not shown). Conversely, induction of additional MTase synthesis decreased the protection, which disappeared completely at or above 1 mM arabinose (Figure 2(e)).

We also monitored the growth of cells carrying the RM system on the pMid plasmid, with or without additional REase or MTase expression plasmids, and control cells carrying empty vectors upon infection with phage λ_vir_ (multiplicity of infection (MOI)=0.2) (Figure 2(f)). Cells were induced with 1 mM arabinose two hours prior to infection. As can be seen, the unprotected culture started to lyse ∼70 minutes post-infection, presumably after the second cycle of phage burst. Cells carrying the RM system plasmid and the pACYC vector continued to grow for ∼150 minutes and then started to lyse, presumably due to accumulation of the modified phage. Induction of additional REase made the culture fully resistant to the phage. Induction of extra MTase synthesis made cells more sensitive to infection compared to cells with the pRM-Mid plasmid as the only source of RM enzymes: lysis started 120 min post-infection and the culture lysed completely by 300 min post-infection when the culture with the pRM-Mid plasmid as the only source of RM enzymes was only partially lysed. Together all these results show that the decrease in MTase to REase ratio increases the level of protection. On the contrary, the increase in MTase to REase ratio decreases the level of protection.

### The influence of PCN and additional expression of REase or MTase on bacteriophage λ DNA degradation after infection of cells carrying the Esp1396I_fluo RM system

We were interested to determine how the copy number of plasmids carrying the Esp1396I_fluo system or additionally expressed individual RM enzymes affect the average lifetime of injected phage DNA inside the cell. To this end, we used the ability of the SeqA protein to specifically bind to Dam methylated or hemimethylated DNA. The host strain for this experiment was *dam-*and constitutively expressed SeqA fused with the mKO2 fluorescent protein (SeqA::mKO2). The DNA of the phage used for infection was fully Dam methylated. Cells carrying pRM-Low, pRM-Mid, or pRM-High plasmids or cells carrying the pRM-Mid plasmids with either +R or +M plasmids supplemented with 1 mM arabinose as well as appropriate control cells were infected with bacteriophage λ_vir_ at the MOI=1. As is described elsewhere (Trinh et al., 2017), SeqA::mKO2 binds to fully methylated injected phage DNA forming a distinct focus visible in the mKO2 channel (Figure 3(a)). When phage DNA is replicated, the number of foci increases to two per each injected phage genome and remains constant throughout the infection until the cell lyses. Accordingly, cells were imaged in transmitted light and in the mKO2 fluorescent channel every 10 minutes after the infection.

**Figure 3.**
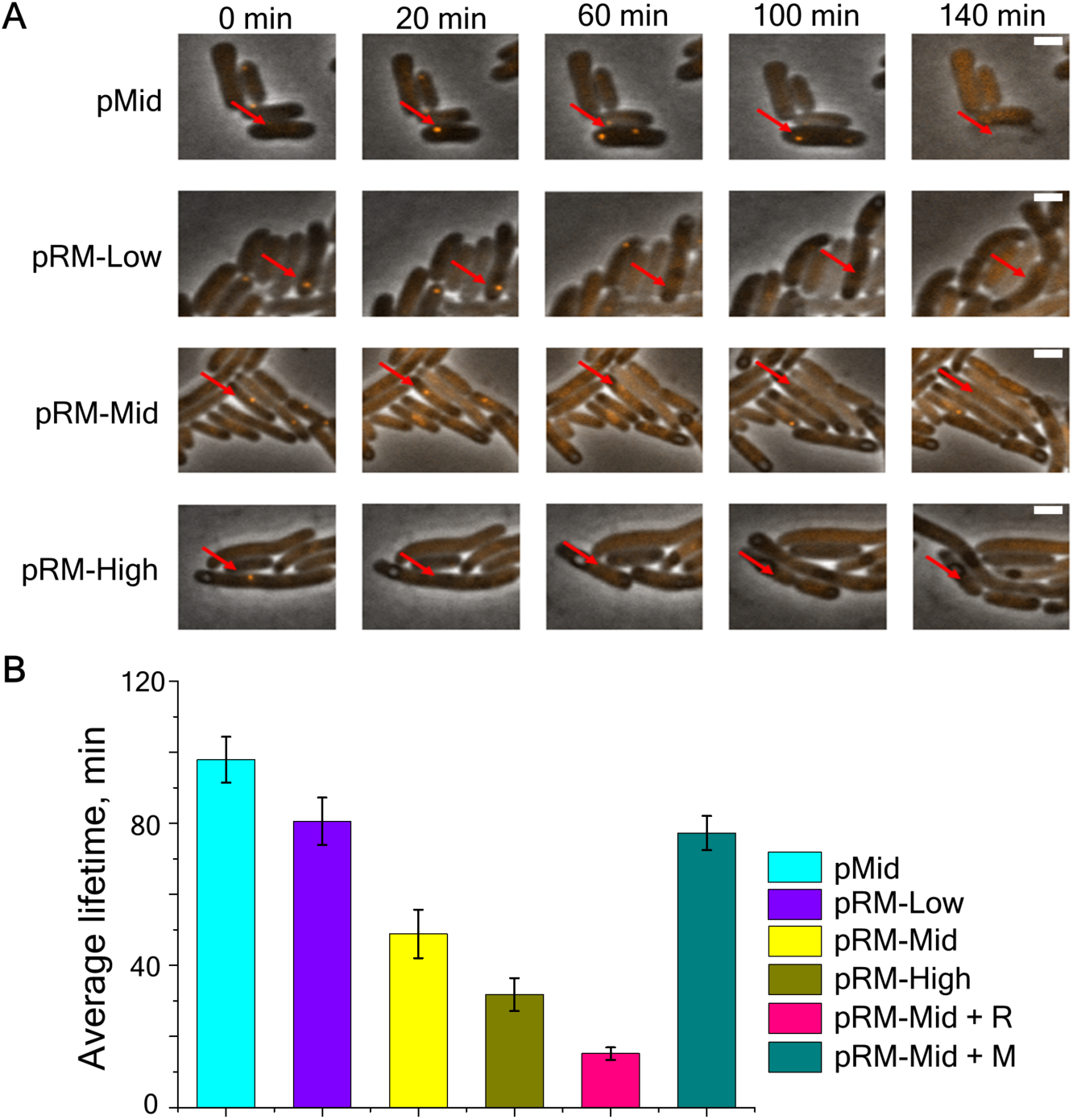
Degradation of phage DNA in single cells protected by Esp1396I. (A) Images from a time-lapse movie showing the appearance of SeqA::mKO2 foci in cells from indicated cultures. Red arrow points to cells infected with phage. Scale bar is 2 μm. (B) Average foci lifetimes in infected cells carrying indicated plasmids. Mean values obtained upon observation of 690 infected cells carrying empty pMid plasmid, 43 cells carrying the pRM-High plasmid, 128 cells carrying the pRM-Mid plasmid, 133 cells carrying the pRM-Low plasmid, 214 induced cells carrying pRM-Mid and +M plasmids, and 49 induced cells carrying the pRM-Mid and +R plasmids are shown. Error bars represent standard errors of the mean.

In pMid cells and in induced pRM-Mid+M cells, the appearance of a single mKO2 focus shortly after the beginning of observation was detected followed by the appearance of the second focus after ∼60 minutes, indicating replication of injected phage DNA, which in turn suggests that the RM protection has been overcome. Indeed, after 140 minutes of observation, such cells lysed. For unprotected pMid cells, an average focus lifetime was 97 minutes (foci lifetimes were calculated from the first moment of appearance of a focus until cell lysis). In bacteria harboring RM plasmids, only single foci were observed, and bacteria with these foci did not lyse; instead the foci eventually disappeared. The average foci lifetimes (measured from the time of focus appearance till its disappearance) in cells harboring pRM-Low, pRM-Mid and pRM-High plasmids were 80, 49 or 31 minutes, respectively. For pRM-Mid samples with additionally induced MTase and REase, these values were, correspondingly, 82 and 18 minutes (Figure 4(b)). We conclude that the presence of an RM system decreases the lifetime of unmodified phage DNA, as expected. Higher concentrations of REase reduce the average time needed for degradation of phage DNA thus enhancing the level of protection. On the contrary, in cells with a higher concentration of MTase, the average time needed for invading phage DNA degradation is increased, leading to a lower level of protection. Increased copy numbers of RM plasmids decrease phage DNA lifetime inside cells, also increasing protection.

**Figure 4.**
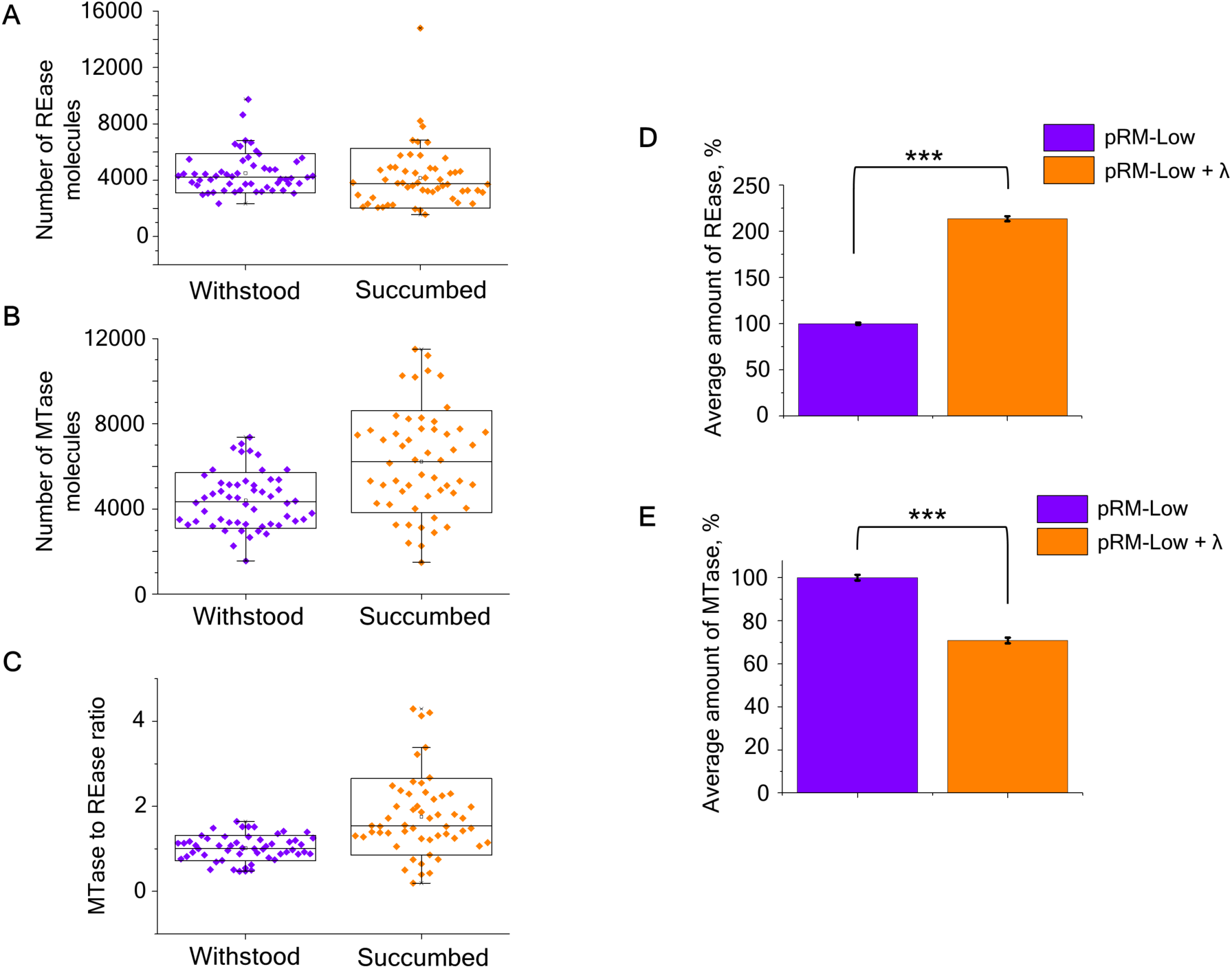
Distribution of enzymes in cells with the pRM-Low plasmid that either succumbed to or withstood phage infection. Distribution of REase (A) and MTase (B) molecule numbers and of MTase to REase ratios (C) in individual productively infected (orange box charts) and withstood (violet box charts) cells carrying the Esp1396I_fluo system on indicated plasmids at the time of the infection. (D) Average amounts of REase in cells carrying the pRM-Low plasmid without (purple) and after (orange) a round of λ_vir_ infection measured by flow cytometry. Average fluorescence in the red fluorescence channel of non-infected cells transformed with the pRM-Low plasmid was taken as 100% of the fluorescent signal. (E) Average amounts of MTase in cells carrying the pRM-Low plasmid without (purple) and after (orange) a round of λ_vir_ infection measured by flow cytometry. Average fluorescence in the green fluorescence channel of non-infected cells transformed with pRM-Low plasmid was taken as 100% of the fluorescent signal.

### Determination of REase and MTase levels in productively infected bacteria

We next determined whether there is a specific subpopulation of cells in RM-protected cultures that is more efficiently attacked by unmodified phage. Based on results presented above, cells in such a subpopulation, if it exists, should have a higher MTase to REase ratio than cells in the rest of the population. We infected bacteria transformed with the pRM-Low plasmid with bacteriophage λ _vir_ and followed the infection using live fluorescence microscopy. The pRM-Low plasmid was used because protection levels afforded by pRM-Mid and pRM-High plasmids were too high, making it impossible to observe enough rare productive infection events under the microscope. Infection was carried out at the MOI of 1. While 33% of cells transformed with the pLow plasmid were lysed after a round of infection, only 7% of cells carrying the pRM-Low plasmid lysed, in agreement with the protection levels determined using the drop test method (Figure 1(c)). Given that the experiment was conducted at high MOI, we surmise that most surviving RM-bearing cells were infected at the beginning of the experiment but managed to destroy injected phage DNA. We estimated the overall levels and the ratios of MTase and REase fusions in 54 RM-carrying cells that succumbed to the infection and in the same number of cells that withstood it (Figure 4(a), (b) and (c)). The REase and MTase amounts were measured at the first frame, taken 10 minutes after infection.

The mean numbers of REase fusion molecules in productively infected cells and cells that withstood the infection were, respectively, 4155±288 and 4500±190 per cell, which is statistically indistinguishable. However, the difference in mean numbers of MTase fusion molecules in infected cells and cells that withstood the infection (6219±326 and 4400±178, respectively) was highly significant (*p* < 0.001) and led to a shift of the MTase to REase ratio in productively infected cells to higher values compared to cells that withstood the infection (1.75±0.123 versus 1.02±0.04).

As an alternative method to investigate whether a round of phage infection changes the MTase to REase ratio distribution in a surviving bacterial population, infected and uninfected cultures of cells carrying the pRM-Low plasmid were analyzed by flow cytometry. Infection was performed at a high MOI=5 such that all cells were expected to be infected at the start of the experiment. A shift in the distribution of fluorescence intensities in cells upon the infection was detected. First, a significant increase (two-sample Kolmogorov-Smirnov test p-value < 2.2*10^-16) in mean fluorescence intensity in the red fluorescence channel corresponding to REase was observed (Figure 4(d)). Second, there was a significant (Kolmogorov-Smirnov test p-value < 2.2*10^-16) decrease (Figure 4(e)) in mean fluorescence intensity in the green fluorescence channel corresponding to MTase. We conclude that cells with stochastically higher concentrations of REase or lower concentrations of MTase become more prevalent in the population of cells with the plasmid-borne RM system after one round of infection with the phage. This observation supports a view that cells with a larger than average amount of MTase and a lower-than-average amount of REase are more likely to be infected by the phage.

## DISCUSSION

In this work, we demonstrate that changing the ratio of type II Esp1396I MTase and REase from the one afforded by its natural homeostatic transcription regulation mechanism (Cesnaviciene et al., 2003, Bogdanova et al., 2009) can either completely abolish protection from viral infection (when the amount of Esp1396I MTase is increased) or make all cells resistant to the infection (when the amount of Esp1396I REase is increased). These results confirm the intuitive expectations and show that the balance of MTase and REase activity is critical for protection from genetic predators. Analysis of changes in protection levels at different induction levels of MTase and REase genes confirms our earlier theoretical considerations (Enikeeva et al., 2010) and shows that the MTase to REase activity ratios determine the level of protection.

It is interesting to compare the observed effects of variation in intracellular REase and MTase amounts *N*_*REase*_ and *N*_*MTase*_ on phage infection with theoretical predictions made in (Enikeeva et al., 2010). Specifically, the equation presented in that work,

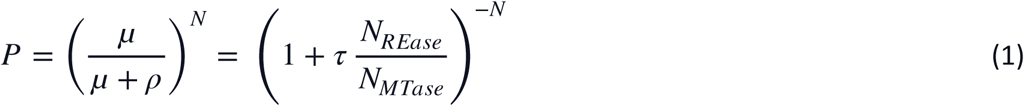

expresses the phage survival probability P through the probabilities of methylation μ and restriction p of all N restriction-methylation sites present in the phage genome. The probabilistic meaning of Eq. (1) is quite transparent: a phage survives and produces (modified) progeny if and only if each of the N sites becomes methylated and no site is cleaved by a REase. The cellular environment is assumed to be uncrowded so that enzymatic reactions at a given recognition site are independent of the fate of other sites. It is further assumed that the concentrations of REase and MTase enzymes are low, such that μ and p are proportional to the corresponding numbers of MTase and REase molecules within the cell, i.e., *μ* = *α*_*MTase*_ *N*_*MTase*_ and *ρ* = *α*_*REase*_ *N*_*REase*_, with factor τ defined as the ratio of proportionality coefficients, *τ* = *α*_*REase*_ /*α*_*MTase*_.

To see how well Eq. (1) describes phage survival probability observed in experiments with cells harboring the Esp1396I_fluo system on plasmids with different PCN or cells expressing additional MTase or REases, we plotted

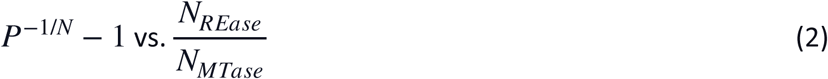

for three data sets: i) data obtained with cells harboring the RM system on high, middle, or low copy number plasmids; ii) data obtained with cells carrying the RM system encoded on a pRM-Mid plasmid with additional expression of MTase from the +M plasmid and grown at different concentrations of arabinose inducer, and iii) data obtained with cells carrying the pRM-Mid plasmid and the +R plasmid grown at different concentrations of inducer. The results are presented in Figure 5. As can be seen, a fit *y* = *τx, τ* = 0.25, expected to be linear from Equation 1, is very good and passes well within the standard deviations of each data set. This indicates that a simple Equation 1 indeed captures the essence of REase and MTase kinetics and predicts phage survival in cells harboring plasmid-borne Esp1396I RM systems. It should be noted that the statistical quality of the pRM-Mid+R dataset (magenta) noticeably differs from that of two other data sets (black and green) (Figure 5) and a significant increase in standard deviation is observed in addition to the expected increase in the REase to MTase ratio values. Microscopic analysis of individual cells indicated that in some cells high levels of expression of the REase gene led to unstable and uneven production of the REase fusion, a toxic protein. A number of particularly bright cells became elongated and presumably underwent a SOS response caused by cleavage of undermethylated genomic DNA.

**Figure 5.**
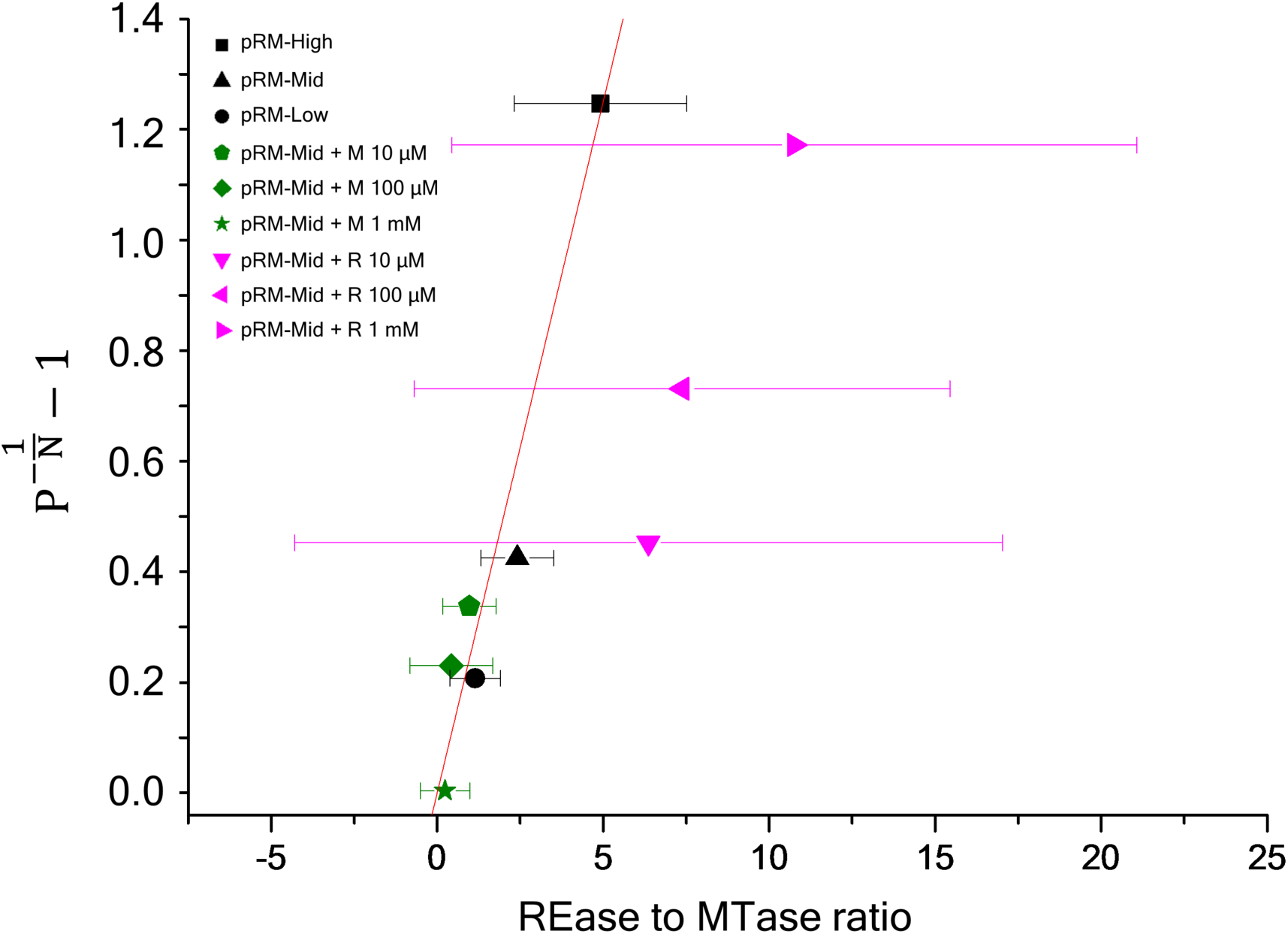
The least square linear fit *y* = *τx* to three datasets shown in black (bacterial cells with pRM-High, pRM-Mid, pRM-Low plasmids), green (pRM-Mid+M - cells with pRM-Mid and +M plasmids in presence of different concentrations of arabinose), and magenta (pRM-Mid+R - cells with pRM-Mid and +R plasmids in presence of different concentrations of arabinose). The fit has the S1ope τ = 0.25 and is shown by a red line. For each dataset N =14 and the respective average and standard deviation are shown by a symbol and horizontal error bars

The actual levels of protection observed in cells with RM systems can vary depending on the system and must be subject to evolutionary pressures. The exact nature of these pressures remains to be defined, though the increased amount of REase needed to achieve a practically complete protection from phage infection likely decreases the viability of cells below a tolerable level by allowing occasional attack of host own DNA before it is fully methylated.

Our demonstration that alterations in the MTase to REase ratio can drastically affect the level of phage protection provides an explanation of how the protection afforded by RM systems can be overcome in clonal cell populations. Stochastic variation in intracellular amounts of RM enzymes may lead to increased susceptibility of individual cells to infection and DNA of progeny phage of such infection events shall be modified, allowing it to take over the initially protected population of cells. Indeed, our real-time analysis of individual infected cells harboring the Esp1396I system on a low copy number plasmid shows that productively infected cells at the time of infection have relatively more MTase than cells that ward off the infection. Thus, the tight regulation of Type II RM systems genes transcription, serves not just to establish a correct temporal sequence of MTase (first) and REase (second) synthesis during the establishment of an RM system-carrying plasmid in a naïve host (Morozova et al., 2016), but also affords the highest possible level of protection by allowing stable and robust levels of expression of both genes.

## ACKNOWLEDGEMENTS

The research is funded by the Ministry of Science and Higher Education of the Russian Federation under the program ‘Priority 2030’ (Agreement 075-15-2021-1333 dated 30.09.2021). Y.I. acknowledges support from FONDECYT project 1200708.

## DATA AVAILABILITY STATEMENT

Materials and reagents described in this article are freely available from the authors upon request. Datasets from Flow Cytometry experiments have been deposited in FlowRepository (Repository ID: FR-FCM-Z55J, http://flowrepository.org/id/FR-FCM-Z55J).

## Supplementary material

**Supplementary Figure S1.**
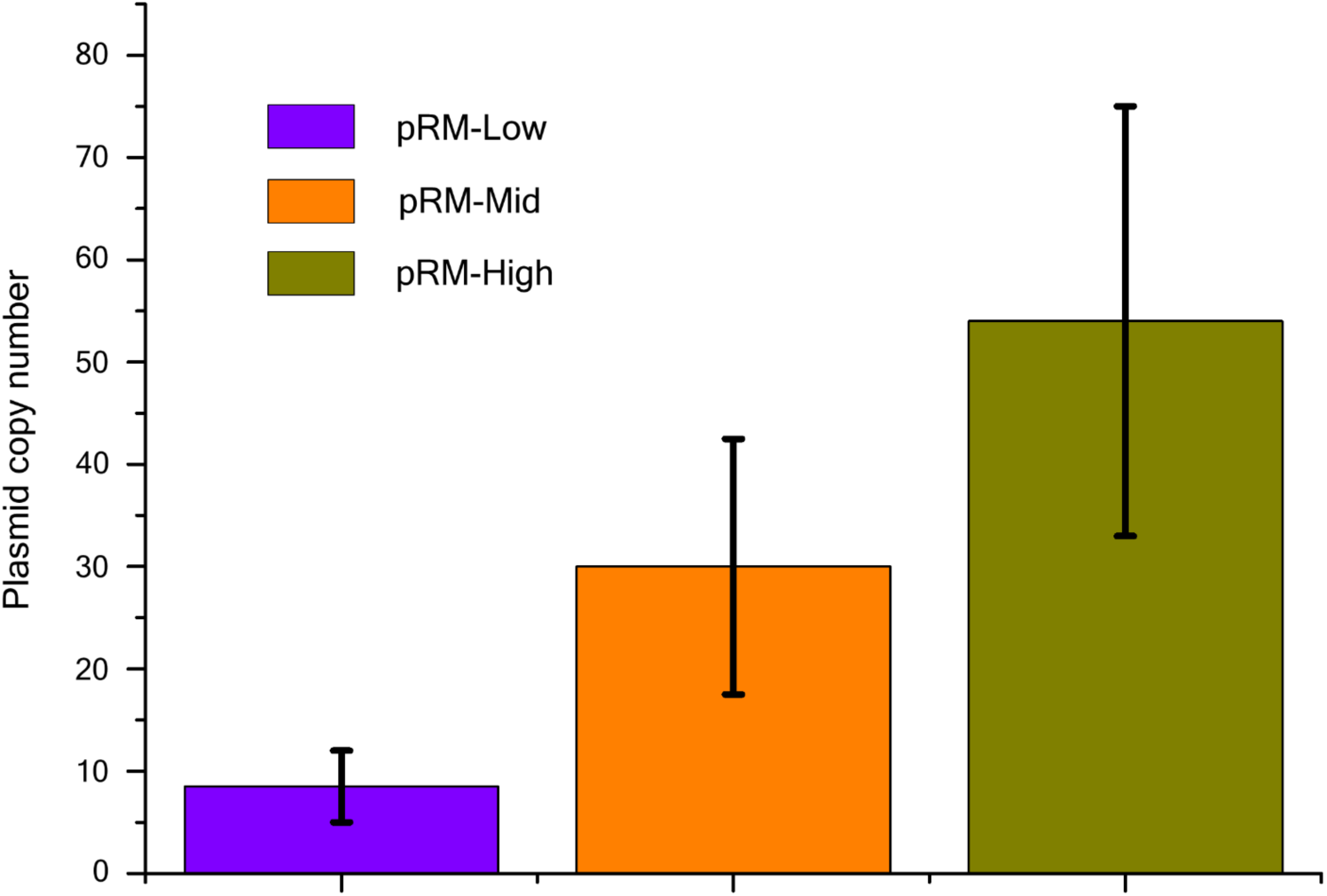
Plasmid copy number (PCN) of pRM-Low, pRM-Mid, and pRM-High plasmids. The PCN was determined using real-time PCR using the ΔC_t_-method. The concentrations of plasmid and chromosomal DNA was determined using two separate primer sets, specific for the Esp1396I methyltransferase gene and the *gyrA* gene (coding for the DNA gyrase subunit) (primer sequences are provided in Table S1). To calculate the absolute concentration of plasmid and genomic amplicons, efficiency of PCR was estimated from standard curves obtained from real-time PCR reactions with aliquots of serial dilutions of solutions containing known concentrations of purified pRM-High or chromosomal DNA as templates. PCN was calculated by taking a ratio of absolute concentrations of plasmid and genomic DNA in the sample. Error bars represent standard deviations and the bars show mean values obtained from measurements performed for three independent cultures.

**Supplementary Figure S2.**
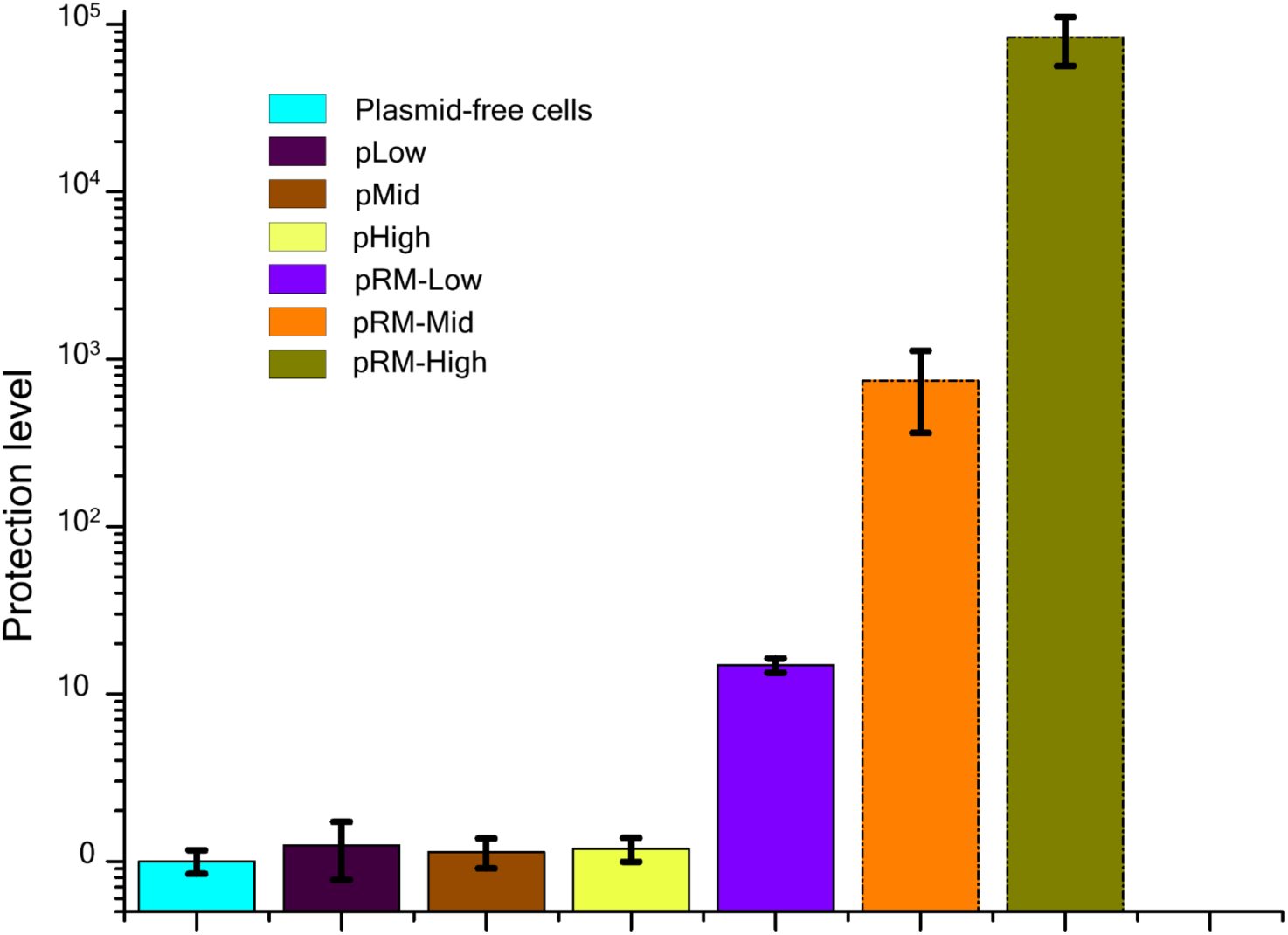
Protection of plasmid-free cells or cells carrying indicated plasmids from phage λ_vir_ infection. Protection levels were calculated by dividing the titer of phage lysate on the lawn of plasmid-free DHS*α* cells by a titer determined on lawns of cells carrying an indicated plasmid. Bars represent mean protection levels obtained from three independent experiments. Error bars show standard errors of the mean.

**Supplementary Figure S3.**
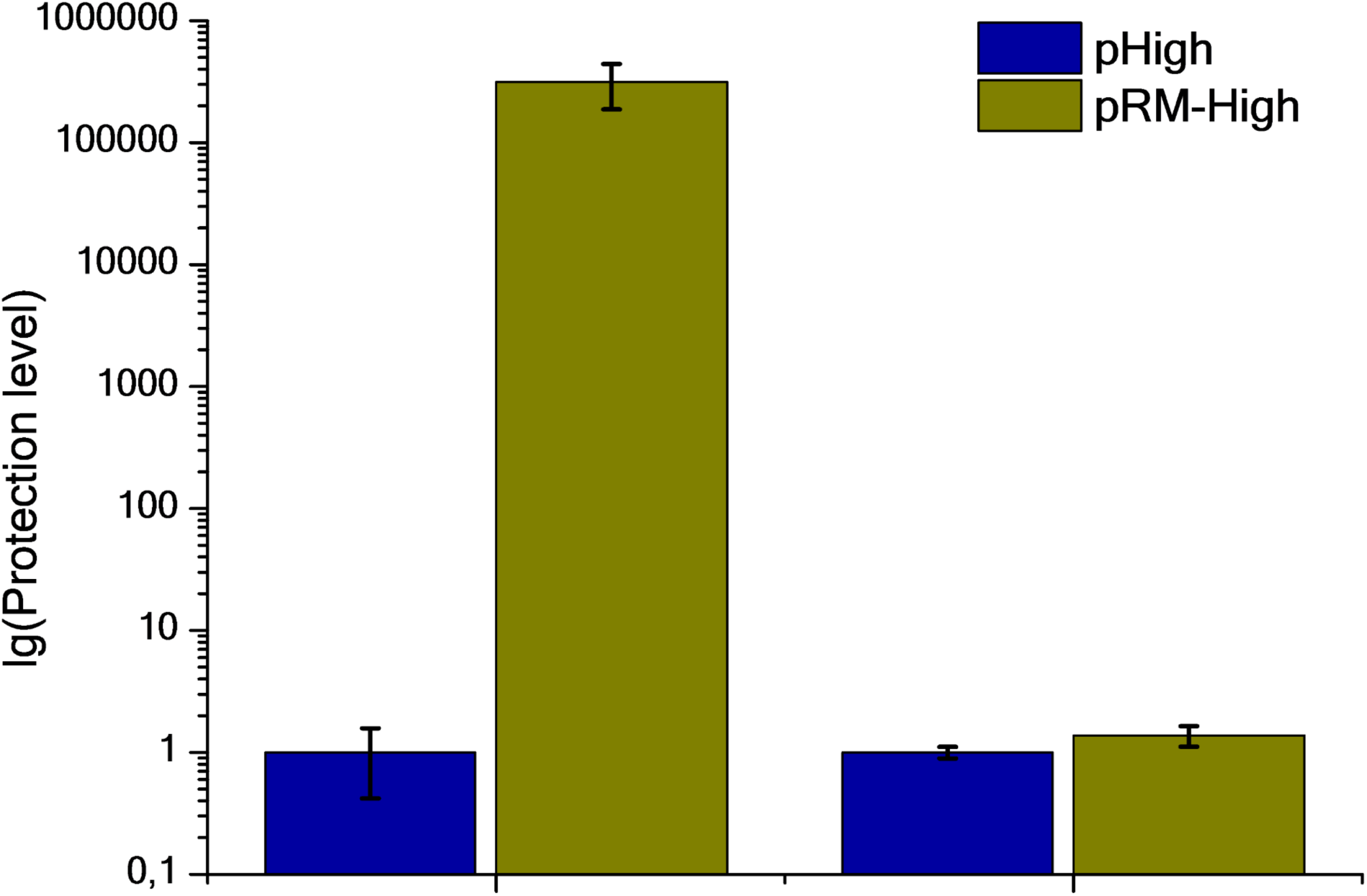
Protection level of cells carrying the pRM-High plasmid or the pHigh vector control from λ_vir_ phage formed after a single round of infection of cells without an RM system (λ_1_) or cells carrying the pRM-Low plasmid (λ_2_). Bars represent mean protection levels obtained from three independent experiments. Error bars represent standard errors of the mean.

**Supplementary Table S1.**
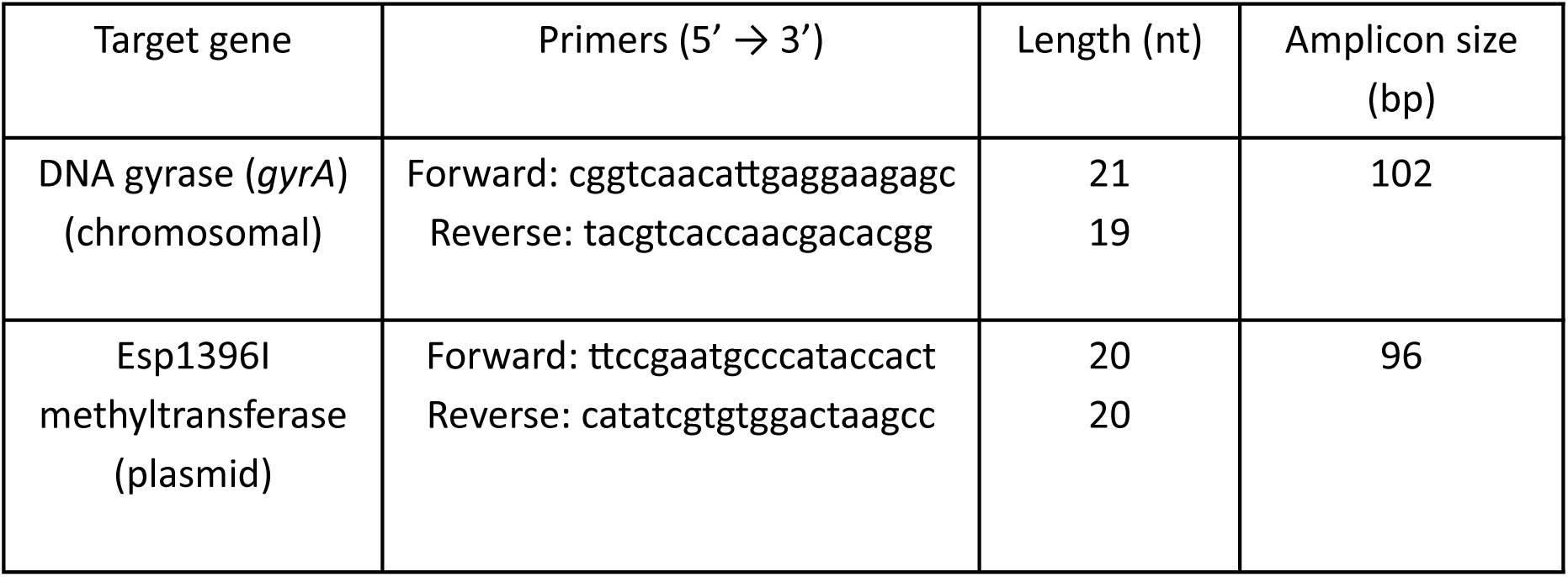
Primers for real-time PCR.

